# RNA polymerase is a molecular motor that converts its transcribing genetic information to free energy for its movement

**DOI:** 10.1101/2023.04.10.532525

**Authors:** Tatsuaki Tsuruyama

## Abstract

RNA polymerase (RNAP) is an enzyme that catalyzes RNA synthesis from a DNA template via translocation on the DNA. Several studies on RNAP translocation have shown an unexplainable discrepancy in the experimental value of the average free energy change (*ΔG*) required for RNAP translocation. To address this inconsistency, we propose a model of the transcription system based on information thermodynamics integrating information theory and thermodynamics. The state function of RNAP was defined from its position on the template DNA, its migration direction, and the deoxyribonucleotide (dNTP) that it transcribes. Based on the state function, *ΔG* was defined consisting of *ΔG*_*th*_ required for determining its position *m* and other thermodynamic factors, and *ΔG*_*d*_ required for determining movement orientation or *ΔG*_*N*_ value from the dNTP sequence information of the DNA transcribed by RNAP. *ΔG*_*d*_ was calculated using the fluctuation theorem applying movement orientation and *ΔG*_*N*_ was estimated based on information thermodynamics of mutual information given by dNTP appearance ratio. It was found that the involvement of either *ΔG(d)* or *ΔG(N)* in free energy caused a discrepancy in *ΔG* values. In conclusion, information thermodynamics can be a framework for information processing in the cell, and RNAP serves as a good model of molecular machine moves through the conversion of information.

**Significance Statement:** RNA polymerase directly converts deoxyribonucleotide sequence information from template DNA to free energy for its movement along the DNA strand.

## Main text

Our understanding of molecular machines from a biophysical viewpoint has recently advanced, with membrane transport (1-3), signal transduction (4, 5), and molecular motors (6-10) such as RNA polymerase (RNAP) being extensively studied (11-19). RNAP catalyzes RNA synthesis by adding ribonucleotides (rNTPs) onto the RNA precursor, i.e., by transcription, from the deoxyribonucleotide (dNTP) of the DNA template. When the transcription of one dNTP is completed, RNAP moves through thermodynamic fluctuations between the 3′→5′ and 5′→3′ orientations along a DNA template strand for the next dNTP. This RNAP movement has been postulated to be “powered” by the chemical energy from rNTP hydrolysis during rNTP addition (20, 21).

Several experimental studies have estimated the average free energy change (Δ*G*) of RNAP during translocation. Thomen *et al*., Wang *et al*., and Guo *et al*. estimated Δ*G* to be −1.3 *k*_*B*_*T* (11, 22), −0.2 to 3.4 *k*_*B*_*T* (23), and 0.0 *k*_*B*_*T* (24), respectively, where *k*_*B*_ is the Boltzmann’s constant and *T* is the temperature of the transcription system. The significant difference (almost 5.0 *k*_*B*_*T*) between these values has puzzled scientists (15, 23) (**Fig. 1**). To address this inconsistency, we assumed that there is an unknown thermodynamic parameter; therefore, an information thermodynamics model of the transcription system is proposed. Here, RNAP may have the ability to transform the information of the deoxyribonucleotide (dNTP) type within the template DNA (e.g., dGTP in **Fig. 1**) into the work of auto-translocation along the template DNA. Among RNAP, T7 RNAPI has been used to measure RNAP movement, and we modeled this enzyme (12, 22, 24-26).

**Fig. 1.**
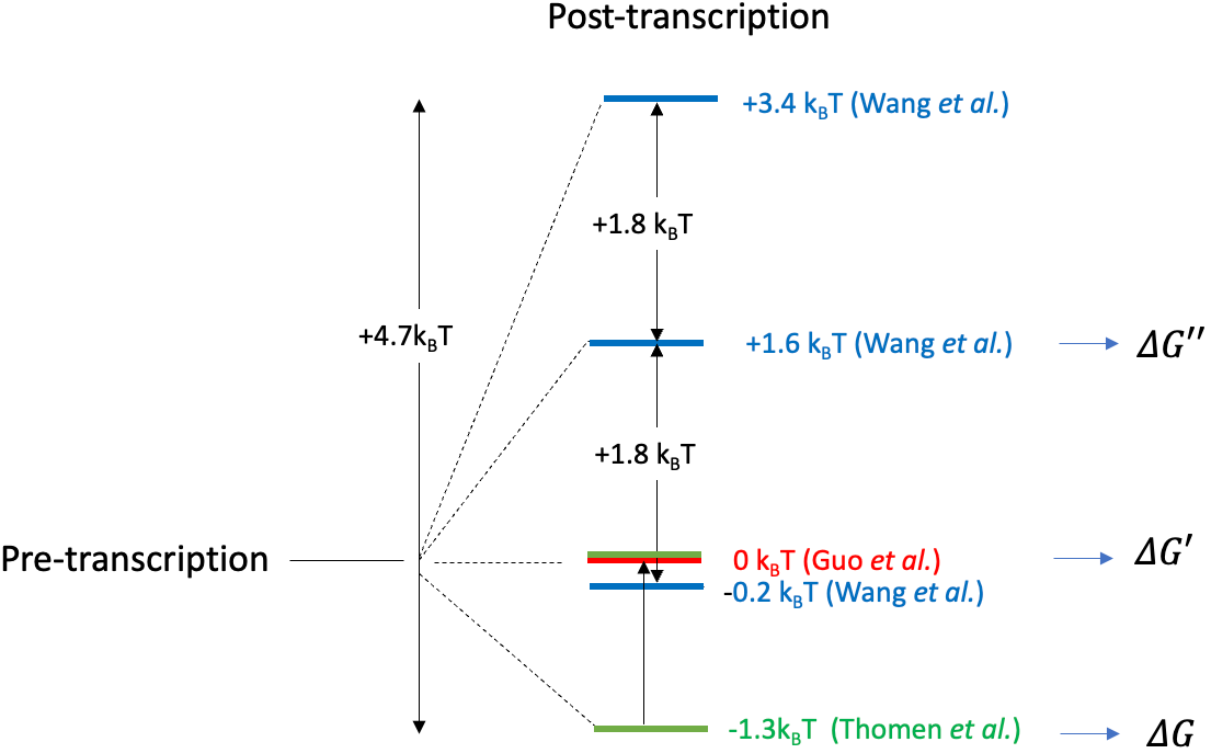
Difference in free energy levels between pre- and post-DNA transcription via RNAP. The colored horizontal lines represent the RNAP energy levels estimated in different studies; the blue lines depict the range of −0.2 to +3.4 *k*_*B*_*T* reported by Wang *et al*. (25); the green line depicts the value reported by Thomen *et al*. (24); the red line depicts the value reported by Guo *et al*. (26). *ΔG, ΔG*′, and *ΔG*′′ indicate the free energy change variations in the text.

## Results

### Transcription system model

We introduced a function in the transcription system model that describes the state of RNAP. This model consists of RNAP, a DNA template strand comprising deoxyribonucleotide, *N* = dNTP (dGTP, dATP, dCTP, dTTP), an RNA precursor consisting of ribonucleotide, *R* = rNTP (rCTP, rUTP, rGTP, rATP), and an rNTP pool for the elongation reaction onto the precursor RNA in the cytoplasm (**Table 1**). Transcription proceeds in four steps: reading the dNTP of the template (here, dGTP) to be transcribed, recruitment of rNTP (here, rCTP) to be incorporated into the transcriptional system (rNTP recruitment), addition of rNTP to the RNA precursor (elongation), and RNAP translocation (**Fig. 2**). During translocation, RNAP moves at a one-nucleotide interval step-by-step, fluctuating between forward (3′→5′) and backward (5′→3′) orientations along the template DNA from the 1^st^ to the *m*^th^ dNTP.

**Table 1.** Experimental RNAP translocation data. *f* denotes the force of RNAP movement. *h* denotes the mutual information gained by RNAP from template DNA. The work done on RNAP by the DNA template is denoted by *W* = *fd/k*_*B*_*T. ΔS/k*_*B*_ is calculated using the ratio *P*^+^(*Δs*_*tot*_*/k*_*B*_)/*P*^−^(−*Δs*_*tot*_*/k*_*B*_) = *t*^+^*/t*^−^ (17, 18, 22).

| $\log \frac{P_+(\Delta S_{tot})}{P_-(-\Delta S_{tot})} (10^{-21} \text{ JK}^{-1})$ | $f (10^{-12} \text{ (N)})$ | $W (10^{-21} \text{ (J)})$ | $\Delta G (10^{-21} \text{ (J)})$ | $-\Delta G + k_B T H (10^{-21} \text{ (J)})$ |
| --- | --- | --- | --- | --- |
| 4.0 (17, 22), 2.8(18) |  |  |  |  |
|  | 10 (15) | 0.82 | -0.57 | 1.95 |
|  | 11 (16) | 0.9 | -0.49 | 1.87 |
|  | 15 (16) | 1.23 | -0.16 | 1.54 |
|  | 18 (17) | 1.48 | 0.09 | 1.29 |
|  | 20 (34) | 1.64 | 0.25 | 1.13 |
|  | 25 (16) | 2.05 | 0.66 | 0.72 |
| Average | 16.5 | 1.35 | -0.04 | 1.42 |

**Fig. 2.**
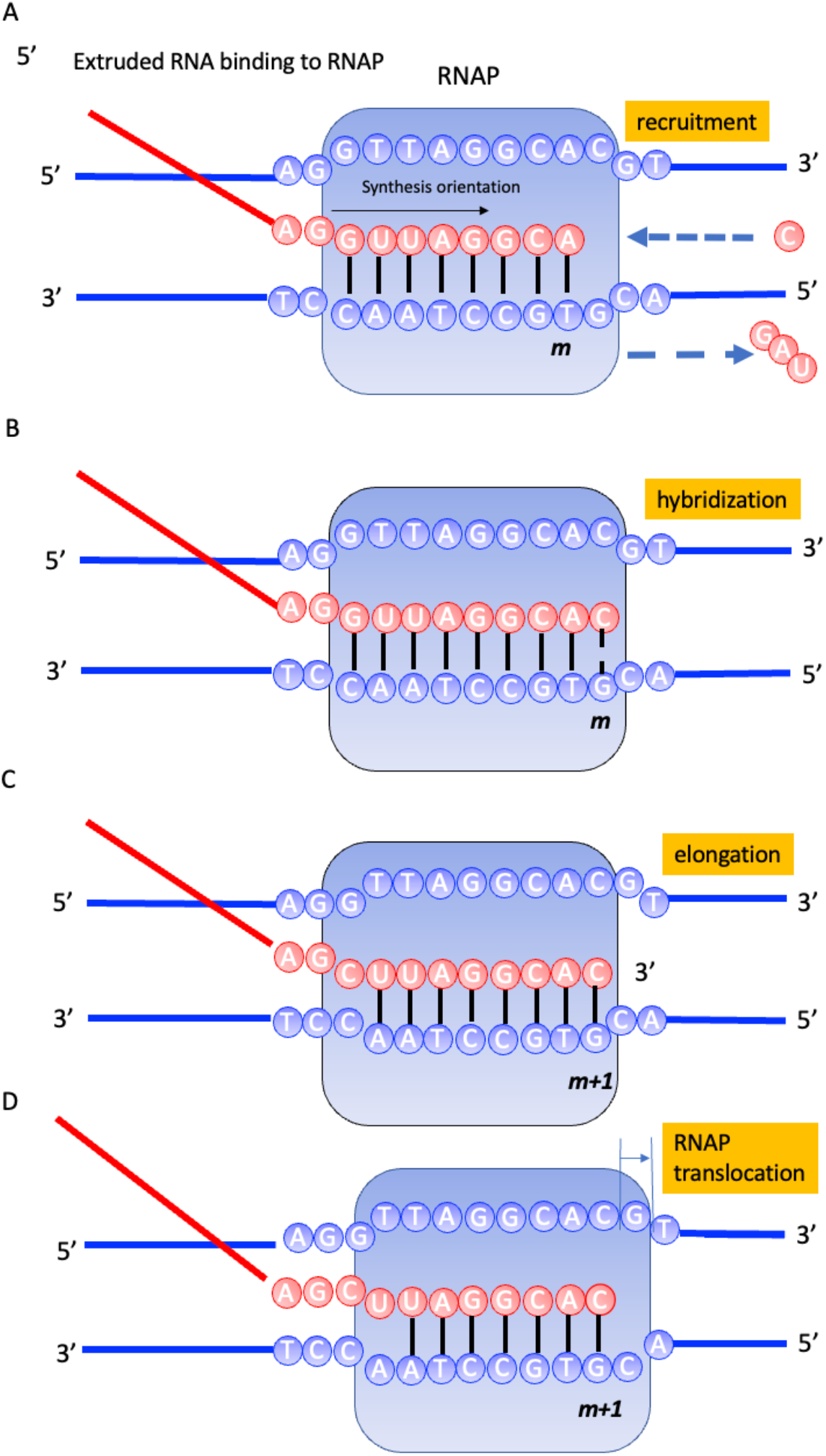
Model of transcriptional action. The four steps of RNA transcription. The blue lines and circles represent template DNA. The red lines and circles represent the RNA precursor. (A) dGTP is being transcribed, rCTP is being recruited, and rGTP, rATP, and rUTP are not being recruited (red circles). (B) Recruited rCTP hybridizes with dGTP (hybridization). (C) Hybridizing rCTP is incorporated at the 3′- end of the RNA (RNA elongation). (D) Translocation of RNAP by one dNTP.

### Stochastic thermodynamics of translocation

The system was supposed to be isobaric and isothermal. Considering the status function of RNAP, the state of RNAP is described by (i) the template DNA binding position on the template DNA (indicated by the coordinate *m*; the position on the DNA template segment between the *m*^th^ dNTP and the (*m*+1)^th^ dNTP), (ii) *d*; movement direction, 3′→5′ (+) or 5′ → 3′(-), and (iii) *N*, the dNTP of the nucleotide of the DNA. Here, we defined the probability density functions *P*(*m*) for (i), *P*(*d*) for (ii), and *P*(*N*) for (iii) and defined the total probability function *p*(*m, d, N*) given by Eq. 1:

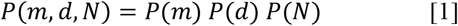

Furthermore, the entropy of the transcriptional system is defined by Eq. [1] using the probability distribution:

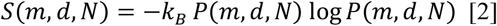

Equation [2] calculates the expected value for − log*p*(*m, d, N*), which we can further define as:

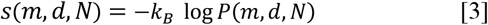

where *s* is the stochastic entropy of RNAP, which is described by the sum of entropy, *s*_*th*_ (*m*) *=* -*k* _*B*_ log*P*(*m*), *s*_*d*_ = - *k*_*B*_ log *P*(*d*), and *s*_*N*_ = - *k*_*B*_ log *P*(*N*):

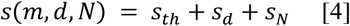

According to the above equations, we defined free energy as Δ*G* = -*T* Δ*s*, Δ*G*_*th*_ = -*T* Δ*s*_*th*_, Δ*G*_*d*_ = -Δ*s*_*d*_, and Δ*G*_*N*_ = - *T* Δ*s*_*N*_. However, in the definition, we neglected internal energy due to isothermality. Below, we introduce a thermodynamic model of template DNA transcription by RNAP.

### The probability function and entropy of RNAP

An example of the RNAP trajectory along the DNA template strand is shown in **Fig. 3**, where RNAP moves via fluctuations between the 3′→5′ and 5′→3′ orientations. In the process of forward RNAP migration, in the first step, RNAP binds to the 3′-end of the target DNA (*m =* 1) with probability *P*(*m =* 1), and then, it moves to *m* →*m*+1 (1 ≤ *m* ≤ *N*) at the transition rate *k*_*m*_+, finally reaching the 5′-end (*m = N*), where RNAP dissociates from the DNA. In the process of RNAP migration in the opposite direction, RNAP is transferred to the 5 ′ -end (*m = N*) with probability *P*(*m = N*), then moves to *m*+1→*m* at the transition rate *k*_*m*_-, finally reaching the 5′-end (*m =1*) (3, 25). Upon reaching the 5′-end, RNAP dissociates from the DNA. In the process of forward RNAP migration, the probability distribution function of RNAP at the *m* position is given by:

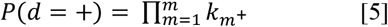

In the process of backward RNAP migration, the probability distribution function of RNAP at the *m* position is given by:

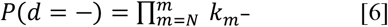

In Eqs [5] and [6],(*d* = +) and (*d* = −) indicate that RNAP moves *m* → *m*+1 for 3′→5′ and *m*+1 → *m* for 5′→3′ in the orientation, respectively. Furthermore, *Q*_*m*+_ (*t*) and *Q*_*m*_*-* (*t*), which represent heat inflow when RNAP moves from *m* → *m*+1 for 3′→5′ and *m*+1 → *m* for 5′→3′, respectively, were introduced. In this case, the thermodynamic entropy change is defined based on Eq. [2]:

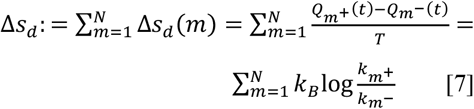

and

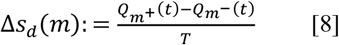

Here, Δ indicates the difference between moving forward and moving backward. Next, we can calculate the entire thermodynamic entropy change as:

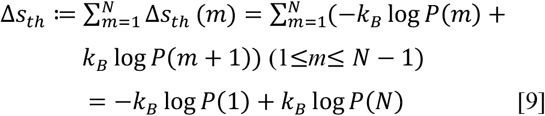

and

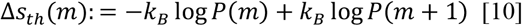

Upon setting the position on the template DNA and considering the movement orientation as an independent factor, Δ*s*_*th*_, the thermodynamic entropy produced in RNAP binding, Δ*s*_*m*_, and movement, Δ*s*_*d*_, can be expressed from Eqs. [7] – [8] as:

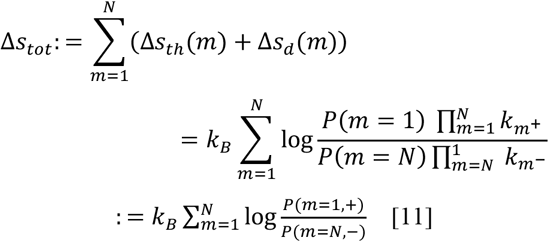

and

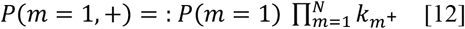

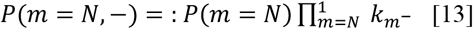

**Fig. 3.**
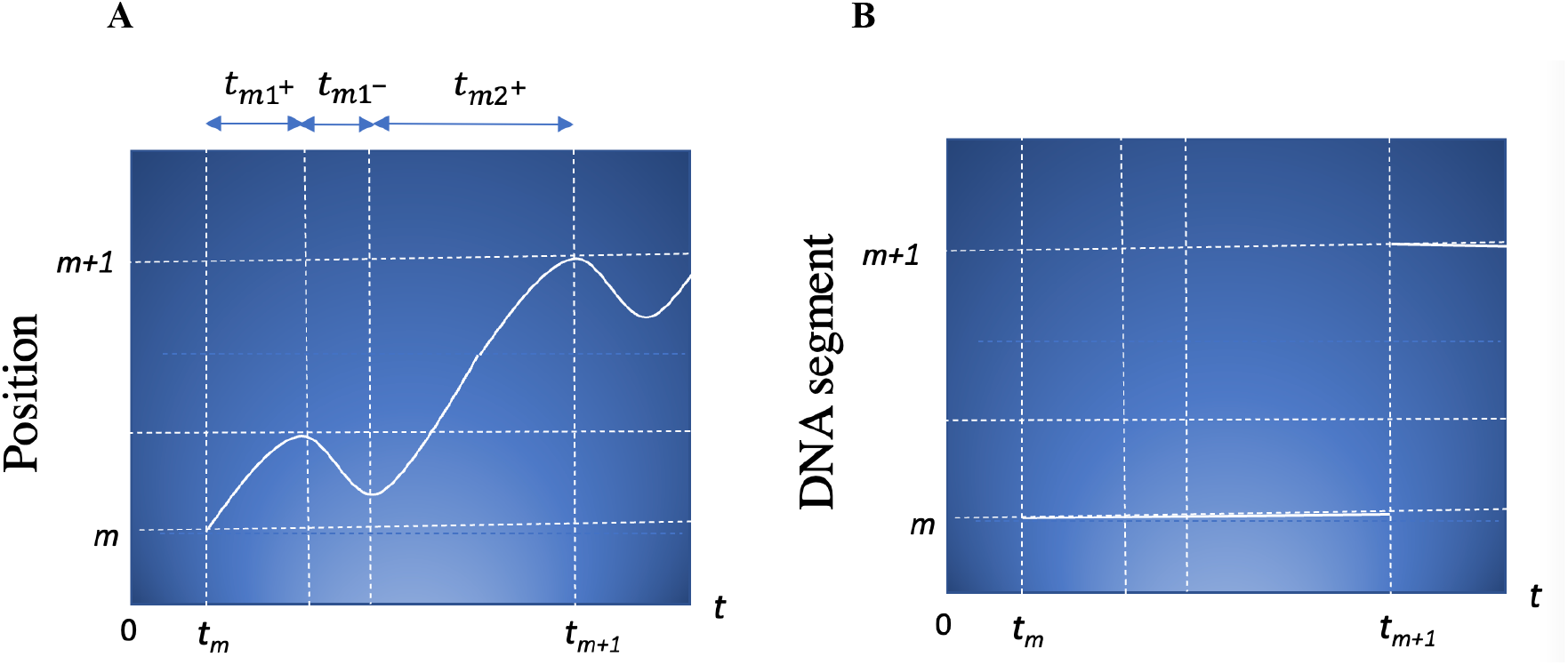
Representative RNAP trajectory along the DNA template strand. RNAP translocation from *m* to *m*+1 fluctuates between the 5′→3′ and 3′→5′ orientations (1 ≤ *m* ≤ *N*). (A) Vertical axes *m* and *m′* (*m*→*m*+1 for 3′→5′, and *m*+1→*m* for 5′→3′) represent the RNAP position of template DNA. (B) Vertical axes represent the DNA segment at which RNAP is located, indicating a discrete value. In both graphs, the horizontal axis *t* denotes translocation time. *t*_*m*_ and *t*_*m*+1_ denote the time when RNAP arrives at the *m* and *m*+1 dNTPs, respectively. *t*_*m1*_^+^, *t*_*m2*_^+^, and *t*_*m1*_^−^ denote the time intervals between *t*_*m*_ and *t*_*m*+1_ when RNAP translocates from 3′→5′ and 5′→3′, respectively. The ratio *P*^+^(*m*)/*P*^−^(*m*) was estimated from *t*_*j*+_/*t*_*j*−_ = ∑_j=1_^2^ *t*_*mj*_^+^/∑_*j* =1_^1^ *t*_*mj*_^−^.

### The path for RNAP movement

Here, we defined the path integrals for each transcriptional orientation of the template DNA. *d*(+) and *d*(–) are additional operations for both RNAP translocation directions, +(*m* → *m+*1) and - (*m*+1 → *m*) respectively (26). From Eq. [11] (**Appendix**):

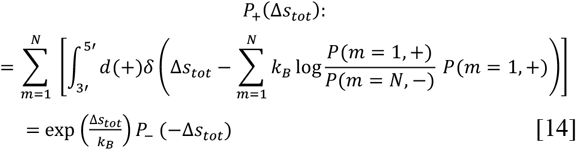

Therefore,

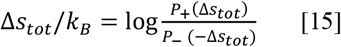

Using the general thermodynamic relation *Δs* = (*W*^*ex*^ *− ΔG*)/*T*, where *ΔG* denotes RNAP *ΔG* without the contribution of the thermodynamic fluctuation and assuming that *W*_*mex*_ (i.e., work done by the template DNA to RNAP) is equal to −*W*, then

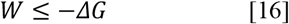

From the experimental data, *W* ∼1.4 *k*_*B*_*T* (Table 1), Eq. [15] was derived:

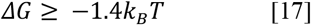

where the obtained lower limit value is approximately equal to the average Gibbs free energy reported by Thomen *et al*. (11, 22). We set:

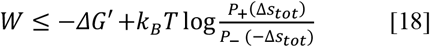

then

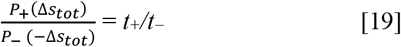

In the above, *t*^+^ and *t*^−^ represent the time necessary for RNAP movement from the 3′-end of the template DNA to the 5′-end and from the 5′-end to the 3′-end, respectively. According to the experimental data, *t*^+^*/t*^−^ ≈ 2.8 - 4.0 (17, 18, 22) (**Fig. 3**), indicating that the movement fluctuates forward and backward by approximately 3.0. Accordingly, log *t*^+^*/t*^−^ = log *P*^+^(-*Δs*_*tot*_)/*P*_*-*_(-*Δs*_*tot*_) = 1.0∼ 1.3 (Table 1). Considering *W* ∼1.4 *k*_*B*_*T* and Eq. [18], Eq. [20] was obtained:

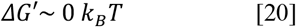

where the obtained value is approximately equal to the lower Gibbs free energy value reported by Wang *et al*. (23) and Guo *et al*. (24).

### Stochastic thermodynamics of reading genetic information

Furthermore, the contribution of the template DNA sequence information was evaluated (**Supplementary Table S1**). In a pre-transcriptional state, the Shannon entropy of *N* is given by *s* (*N*) = −log*P*(*N*), where *P*(*N*) denotes the frequency of each type of deoxyribonucleotide (12). When the transcription error is negligible, the conditional entropy when type *N* is transcribed into *R* by the RNAP is given by *s* (*R*|*N*) = −log *P*(*N*) =−log *P*(*R*). Here, the mutual information in RNAP recognition of *N* as *R* is given by *h* (*R*; *N*): = *s*(*R* − *s* (*R*|*N*) = −log *P*(*R*) – [*−*log *P*(*N*)] = 0. As a post-transcriptional state, *N* does not change, and *R* has already been determined. Therefore, the conditional Shannon entropy is given by *s* (*R* |*N*) = 0. At this point, mutual information *h* (*R, N*) is given by *h* (*R, N*) = *s*(*R*) – *s* (*R* |*N*) = −log *P*(*R*) − 0 = *s*(*R*). Mutual information −log *P*(*R*) is fed back to the template DNA, and *s*(*R*) becomes zero when *R* is determined (**Supplementary Table S2**). In this case, work can be expressed as *ΔG* using mutual entropy *h*(*R, N*) = *s*(*N*), thermal fluctuation, and newly introduced free energy change *ΔG*′′ as follows (5, 27, 28):

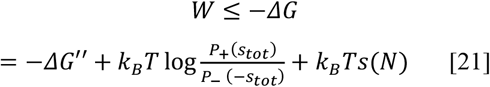

Accordingly, from Eq. [20] and by substituting *s*(*N*) ∼ 1.4 (**Supplementary Table S1**) into Eq. [22], we obtain:

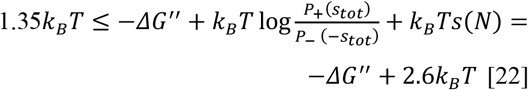

and therefore

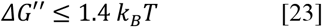

where the obtained lower limit value approximately matches the mean Gibbs free energy value of 1.6 *k*_*B*_*T* reported by Wang *et al*. (23) (**Fig. 1**).

## Discussion

By defining the state function of RNAP, we attempted to determine the information thermodynamics of the DNA transcription process. We found that the discrepancy in the values of free energy changes in enzyme transfer stems from different definitions of free energy. Specifically, free energy takes various values based on whether fluctuations in the direction of movement and the contribution of mutual information are included. In Eq. (21), the free energy is associated with the work done on RNAP. The fact that mutual information can contribute to work is discussed in detail in information thermodynamics, and this is not limited to a specific biophysical model but widely holds.

Several experiments support the conversion of mutual information to physical work (29, 30). The definition of mutual information is theoretically arbitrary and depends on the model, but genetic information is the most essential information for living organisms, and the definition based on the information is essential for understanding biophysical work in the cell.

From the current model, RNAP can be thought of as Maxwell’s daemon that can transform the amount of mutual information into work. The information of the dNTP type of the template DNA to be transcribed is fed back to the DNA by RNAP, preventing RNAP fluctuation along the trajectory of the DNA by anchoring the hybridization of the incorporated rNTP and transcribed dNTP. As a result, RNAP is moved forward predominantly in 3′→5′ direction, indicating work done to RNAP by DNA. This process is similar to the translocation of Brownian particles under the periodic potential of the information-thermodynamic work conversion model in the experiment (30), in which a daemon can move a particle against its potential by repeating the process of obtaining information about the direction of movement of the particle and not returning to the back of the particle (24).

The conversion of genetic information into the translocation mechanism reminds us of the Szilard engine, a refinement of Maxwell’s daemon that can convert particle position information into expansion work by the particle (31). In the model, the RNAP daemon reads the dNTP type of nucleic acid in the template or whether the recruited rNTP is complementary to the template DNA. When rNTP is complementary, RNAP recruits it to the active catalytic center and catalyzes its binding to RNA. In contrast, if rNTP is not complementary, it will not be recruited (32). The obtained information in this selection contributes to the physical migration work of RNAP itself along the template.

In template DNA transcription, RNAP-transcribed information is retained in RNA molecules. Landauer argued that when information obtained from the measurement is stored in the memory (daemon), it is necessary for the daemon to erase the information in the subsequent cycle formation of the step. This argument has widely been accepted as a solution to Maxwell’s daemon paradox. However, because RNAP retains the genetic information as RNA, it is not necessary to erase the memory according to Landauer’s principle. In this way, RNAP can repeat the cycle of reading dNTP information and convert it into self-moving work, which is a molecular motor that can convert thermal fluctuations into work using genetic information. Besides, the current model claims that RNAP possesses the unique property of being able to move itself owing to its own work. This point seems to violate the second law of thermodynamics, and RNAP looks like a molecular machine that cannot be realized unless it can convert the work of mutual information. The fact that RNAP has developed such a mechanism is interesting in terms of molecular evolution.

A limitation of this model is that dNTP type and direction of movement were considered independent factors in Eq. [11]. Frequent fluctuations between at least 3′→5′ and 5′→3′ are approximately 3:1 to 4:1 and may be influenced by the fact that there are four types of nucleic acid nucleotides. It has been experimentally reported that there is a difference in the free energy required for movement for each type (24, 33, 34). Therefore, our model must be validated through experimental verification of free energy for movement by more precise measurements.

In conclusion, the transcription complex represents a unique system capable of directly converting genetic code to thermodynamic work, i.e., its own movement by feedback from template DNA, thereby providing a new perspective on transcriptional regulation and information thermodynamics.

## Appendix

Eq.[12] was given as follow (26):

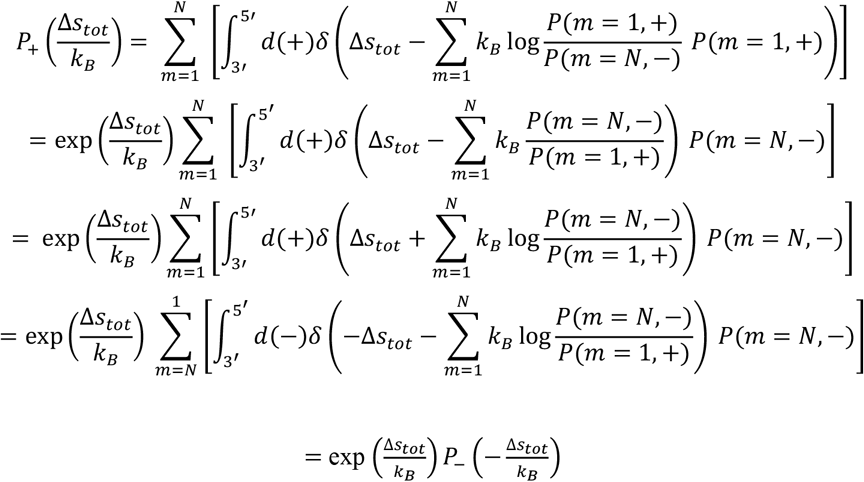

## Acknowledgments

This study was supported by a Grant-in-Aid from the Ministry of Education, Culture, Sports, Science, and Technology of Japan (Synergy of Fluctuation and Structure: Quest for Universal Laws in Non-Equilibrium Systems, P2013-201).

## Data and materials availability

All datasets used in the present study are available in the reports listed in the references.

---

**Table 1.** Abbreviations used in the text.

|  |  |
| --- | --- |
| RNAP | RNA polymerase |
| dNTP | deoxyribonucleotide |
| rNTP | ribonucleotide |
| dNTP | deoxyribonucleotide |
| dGTP | deoxyguanosine triphosphate |
| dATP | deoxyadenosine triphosphate |
| dCTP | deoxycytidine triphosphate |
| dTTP | deoxythymine triphosphate |
| rNTP | ribonucleotide |
| rGTP | guanosine triphosphate |
| dCTP | cytidine triphosphate |
| rUTP | uridine triphosphate |
| rATP | adenosine triphosphate, |
| *m* | position on the DNA template |
| *N* | Deoxyribonucleotide type |
| *R* | ribonucleotide type complementary to *N*<br>(transcribed from *N*) |

## Supplementary Materials

**Supplementary Table 1.** Frequency of each mononucleotide in the T7 promoter of **λ** phage

|  | G | A | T | C | Total |
| --- | --- | --- | --- | --- | --- |
| Probability $P(n)$ | 0.22 | 0.38 | 0.16 | 0.25 | 1.00 |
| $-P(n) \log P(n)$ | 0.33 | 0.37 | 0.29 | 0.35 | 1.33 |

**Supplementary Table 2.**
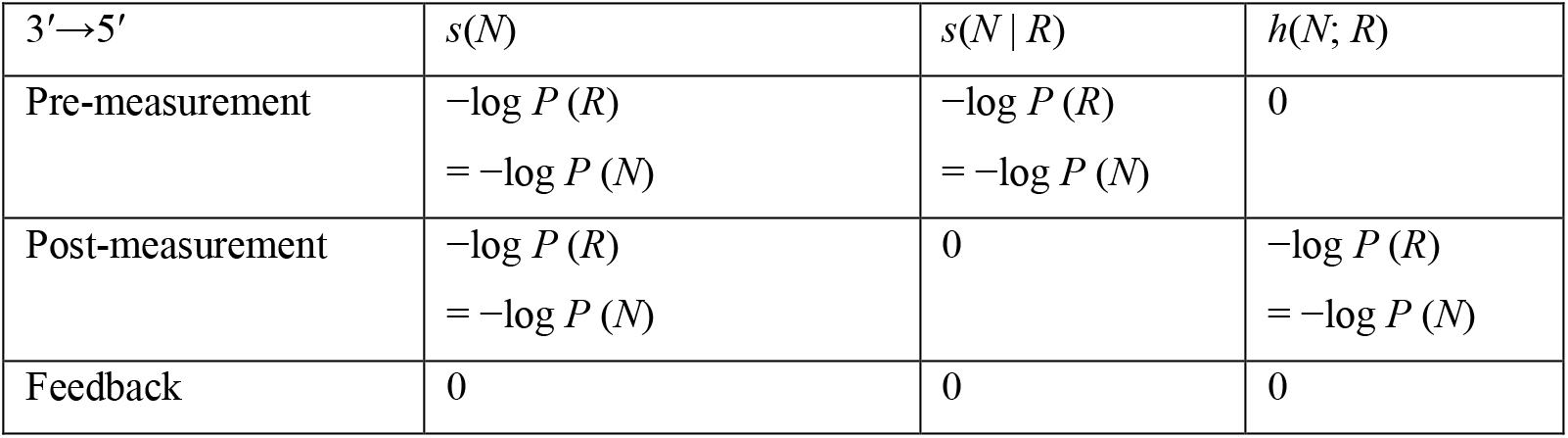
Mutual and information entropy

## Notes

**Competing Interest Statement:** The author has no conflicts of interest to declare.

### Competing Interest Statement

The authors have declared no competing interest.

## References

1. A. M. Berezhkovskii, S. M. Bezrukov, Counting translocations of strongly repelling particles through single channels: fluctuation theorem for membrane transport. Phys Rev Lett 100, 038104 (2008).

2. Y. Hasegawa, Quantum Thermodynamic Uncertainty Relation for Continuous Measurement. Phys Rev Lett 125, 050601 (2020).

3. T. Van Vu, Y. Hasegawa, Geometrical Bounds of the Irreversibility in Markovian Systems. Phys Rev Lett 126, 010601 (2021).

4. T. Sagawa et al., Single-cell E. coli response to an instantaneously applied chemotactic signal. Biophys J 107, 730–739 (2014).

5. S. Ito, T. Sagawa, Maxwell’s demon in biochemical signal transduction with feedback loop. Nat Commun 6, 7498 (2015).

6. D. Andrieux, P. Gaspard, Fluctuation theorems and the nonequilibrium thermodynamics of molecular motors. Phys Rev E Stat Nonlin Soft Matter Phys 74, 011906 (2006).

7. K. Hayashi, H. Ueno, R. Iino, H. Noji, Fluctuation theorem applied to F1-ATPase. Phys Rev Lett 104, 218103 (2010).

8. A. W. Lau, D. Lacoste, K. Mallick, Nonequilibrium fluctuations and mechanochemical couplings of a molecular motor. Phys Rev Lett 99, 158102 (2007).

9. D. G. McMillan, R. Watanabe, H. Ueno, G. M. Cook, H. Noji, Biophysical Characterization of a Thermoalkaliphilic Molecular Motor with a High Stepping Torque Gives Insight into Evolutionary ATP Synthase Adaptation. J Biol Chem 291, 23965–23977 (2016).

10. U. Seifert, Stochastic thermodynamics of single enzymes and molecular motors. The European physical journal. E, Soft matter 34, 1–11 (2011).

11. P. Thomen, P. Lopez, F. Heslot, Unravelling the Mechanism of RNA-Polymerase. Physical Review Letters 94, 128102 (2006).

12. J. Yu, G. Oster, A small post-translocation energy bias aids nucleotide selection in T7 RNA polymerase transcription. Biophys J 102, 532–541 (2012).

13. T. E. Hong-Yun Wang, Alexander Mogilner, and George Oster, Force Generation in RNA Polymerase. Biophysical J 74 (1998).

14. L. Bai, A. Shundrovsky, M. D. Wang, Sequence-dependent kinetic model for transcription elongation by RNA polymerase. J Mol Biol 344, 335–349 (2004).

15. L. Bai, R. M. Fulbright, M. D. Wang, Mechanochemical kinetics of transcription elongation. Phys Rev Lett 98, 068103 (2007).

16. T. W. Turowski et al., Nascent Transcript Folding Plays a Major Role in Determining RNA Polymerase Elongation Rates. Mol Cell 79, 488–503 e411 (2020).

17. E. A. Abbondanzieri, W. J. Greenleaf, J. W. Shaevitz, R. Landick, S. M. Block, Direct observation of basepair stepping by RNA polymerase. Nature 438, 460–465 (2005).

18. D. A. Silva et al., Millisecond dynamics of RNA polymerase II translocation at atomic resolution. Proc Natl Acad Sci U S A 111, 7665–7670 (2014).

19. J. Ma et al., Transcription factor regulation of RNA polymerase’s torque generation capacity. Proc Natl Acad Sci U S A 116, 2583–2588 (2019).

20. C. A. Minetti et al., The thermodynamics of template-directed DNA synthesis: base insertion and extension enthalpies. Proc Natl Acad Sci U S A 100, 14719–14724 (2003).

21. K. S. Dickson, C. M. Burns, J. P. Richardson, Determination of the free-energy change for repair of a DNA phosphodiester bond. J Biol Chem 275, 15828–15831 (2000).

22. P. Thomen et al., T7 RNA polymerase studied by force measurements varying cofactor concentration. Biophys J 95, 2423–2433 (2008).

23. M. D. Wang et al., Force and velocity measured for single molecules of RNA polymerase. Science 282, 902–907 (1998).

24. Q. Guo, R. Sousa, Translocation by T7 RNA polymerase: a sensitively poised Brownian ratchet. J Mol Biol 358, 241–254 (2006).

25. T. Van Vu, Y. Hasegawa, Diffusion-dynamics laws in stochastic reaction networks. Phys Rev E 99, 012416 (2019).

26. Y. Hasegawa, T. Van Vu, Fluctuation Theorem Uncertainty Relation. Phys Rev Lett 123, 110602 (2019).

27. T. Sagawa, M. Ueda, Nonequilibrium thermodynamics of feedback control. Phys Rev E Stat Nonlin Soft Matter Phys 85, 021104 (2012).

28. T. Sagawa, M. Ueda, Fluctuation theorem with information exchange: role of correlations in stochastic thermodynamics. Phys Rev Lett 109, 180602 (2012).

29. T. M. Hoang et al., Experimental Test of the Differential Fluctuation Theorem and a Generalized Jarzynski Equality for Arbitrary Initial States. Phys Rev Lett 120, 080602 (2018).

30. S. Toyabe, T. Sagawa, M. Ueda, E. Muneyuki, M. Sano, Experimental demonstration of information-to-energy conversion and validation of the generalized Jarzynski equality. Nature Physics 6, 988–992 (2010).

31. L. Szilard, On the decrease of entropy in a thermodynamic system by the intervention of intelligent beings. Behav Sci 9, 301–310 (1964).

32. S. Wu, J. Wang, X. Pu, L. Li, Q. Li, T7 RNA Polymerase Discriminates Correct and Incorrect Nucleoside Triphosphates by Free Energy. Biophys J 114, 1755–1761 (2018).

33. L. T. Da et al., T7 RNA polymerase translocation is facilitated by a helix opening on the fingers domain that may also prevent backtracking. Nucleic Acids Res 45, 7909–7921 (2017).

34. J. Ma, L. Bai, M. D. Wang, Transcription under torsion. Science 340, 1580–1583 (2013).

